# Extracting Intercellular Signaling Network of Cancer Tissues using Ligand-Receptor Expression Patterns from Whole-tumor and Singlecell Transcriptomes

**DOI:** 10.1101/127167

**Authors:** Joseph X. Zhou, Roberto Taramelli, Edoardo Pedrini, Theo Knijnenburg, Sui Huang

## Abstract

Many behaviors of cancer, such as progression, metastasis and drug resistance etc., cannot be fully understood by genetic mutations or intracellular signaling alone. Instead, they are emergent properties of cell community that forms a tumor. Studies of tumor heterogeneity reveal that many cancer behaviors critically depend on the intercellular communication mediated by secreted signaling ligands and their cognate receptors that take place between cancer cells and stromal cells. Owing to systematic cancer omics efforts, we studied such cell-cell interactions using data from cancer transcriptome database. We curated a list of more than 2,500 ligand-receptor pairs and developed a method to identify their enrichment in tumors from TCGA pancancer data and to build a cell interaction network from single-cell data for the case of melanoma. Using the specificity of the ligand-receptor interaction and their expressions measured in individual cells, we built a map of a cell-cell communication network which indicate what signal is exchanged between which cell types. Such networks establish a new formal phenotypes which are embodied by the cell communication structure - it may offer new opportunities to identify the molecular signatures which may influence cancer cell behaviors by changing cell population dynamics.

## INTRODUCTION

The burgeoning notion of cellular heterogeneity within a tumor has attracted much attention in cancer research^1,2^. In addition to genetic heterogeneity embodied by the diversity of genomes in each cancer cell due to genome instability^3–5^, it is increasingly recognized that the nongenetic variability of cell phenotypes within an isogenic (clonal) cell population contributes to functional heterogeneity of cancer cells^6,7^. Even not considering the variety from stromal cells (endothelium, stromal fibroblasts and immune cells) but just within tumor, the neoplastic cells raise the possibility of functional specialization and cooperation among the cancer cells themselves^8–11^. Slow proliferating cancer stem cells vs. fast proliferating vs. migratory cells are examples of (overlapping and dynamic) subsets of cells among the neoplastic cells^12,13^. Cooperation would require cell-cell communication which is mainly mediated by physical cellcell contact, shared extracellular-matrix and secreted diffusible signaling molecules^14–16^. Here we study crosstalk between cells via secreted (soluble or matrix) proteins and cell surface signaling factors that bind specifically to their cognate receptors. Herein the principle of autocrine stimulation in which a cell secrete a growth factor and also expresses its cognate receptor has long been used to explain the observed co-expression of both ligand and receptor in the tumor cell population, which was originally considered to be homogeneous and consisting of only one cell type^17^. But the notion of the diversity of cell subtypes among the tumor parenchyma cells which can differ in their capacity to secrete different sets of mediators and to express distinct receptors for these signaling molecules by necessity entails that networks of heterotypic paracrine interactions arise in addition to homotypic interactions between the cells capable of autocrine stimulation.

Studies of heterotypic cell-cell communication mediated by secreted factors and of their role in tumor progression have largely focused on the tumor-stroma cooperation^18–20^. For instance, IL8 secreted by many carcinoma attracts tumor promoting macrophages^21^, and secretion of VEGF stimulates endothelial migration and proliferation, fostering tumor angiogenesis^22^. But with the growing awareness of heterogeneity of the tumor-parenchyma proper, attention has also been given to cooperation amongst distinct cancer cell subpopulations^23,24,9^. For instance, in a study of small cell lung cancer in animal models, the grafting of tumor cell clones genetically engineered to produce different cytokines revealed the effect of cooperative interactions between distinct (engineered) clones of cancer cells on disease progression and metastatic processes. Importantly, in such work, implantation of individual clones separately could not exert any detectable effect^25^.

Similar findings were obtained with a *WNT1*-signaling-dependent breast cancer model where cooperation between two different differentiation lineages (luminal and basal) were required for tumor growth^26^. In a p53-deficient mouse breast cancer model, a crosstalk between tumor initiating cells and a more differentiated mesenchymal cell population was observed^9^. In this study, interaction between different cell populations (defined by different levels of *CD29* and *CD24* markers, respectively) was documented by the detection of secreted ligands, such as *WNT9A, WNT2, IL-6* and *CXCL12* from one cell population which induced a response in the other population through the respective cognate receptors. Furthermore, cytokines produced by the more differentiated mesenchymal cell population were able to stimulate the self-renewal and tumor initiating capacity of the tumor initiating cells. In another case based on the prostate cancer PC3 cell line, a non-cancer stem cell subpopulation rendered a cancer cell subpopulation metastasis-prone^27^. Here, a paracrine interaction between two clonally distinct subpopulations mediated by several diffusible factors, among which protein *SPARC* had a prominent role, resulted in enhanced invasiveness and metastatic dissemination of the cancer stem cell-rich subpopulation of the PC-3 cells.

As described in the paper of Marusyk et. al.^11^ and more generally in the review papers of cytokines and chemokines^28–32^, a large set of growth factors and cytokines/chemokines along with their cognate receptors mediate the cell-cell interactions between different cells in the cancer tissue or even between sub-clones among the cancer cells. Thus, in addition to crosstalk with the non-neoplastic stroma, a large array of secreted proteins and their receptors are produced by the cancer cells themselves for other cancer cells within a tumor. Given the wide functional variety of cells among the tumor cells, we should consider that a substantial promotion of such intra-tumor cell-cell signaling consists of heterotypic interactions whereby some cells send signals designated only for another subset of cells, perhaps only minimally distinct, that expresses the respective cognate receptors.

Since communication is mediated by specific ligand-receptor interactions information about the set of specific “sender-receiver cell pairs” will allow one to define an entire cell-cell communication network in the cancerous tissue in which both neoplastic and stromal cells participate. This network is “hardwired” by the distribution of the cell identities that also dictate the expression of ligand and receptors. Mapping out this communication network would require single-cell resolution gene expression analysis to determine which cell type expresses which ligand or receptor. Such data are, despite the arrival of single-cell transcriptome analysis^33^ still scant.

Currently, the databases of expression of ligands and receptors are derived from bulk tumor (whole tissue) analysis of gene expression profiles. To evaluate if such aggregate measurements, which lack single-cell level information but are readily available for a large number of tumors, provides information on cell-cell communication, we searched for a relationship in the expression of cyto/chemokines and their specific cognate receptors in tumor tissues at the transcription level in the transcriptome data of The Cancer Genome Atlas (TCGA) project^4^. The rationale was that a difference in the expression levels and/or in the correlation of expression of receptors and their ligands between the tumor and the corresponding normal tissues may offer an indication for the role of specific cell-cell communication that is augmented/altered in neoplastic compared to non-neoplastic tissue. The analysis of a total of 5,362 transcriptomes from ten different solid tumors and comparison with corresponding normal tissues revealed a significant alteration of the expression levels and of the correlation of cyto/chemokine transcript abundances with that of their cognate receptors in the tumor tissues, suggesting possible shifts in the communication network associated with malignancy. This finding suggests a role for the non-cell autonomous processes in tumor growth. Constructing cell-cell communication network from whole-tumor tissue profiles may also help in designing new therapeutic strategies that target not intracellular networks - which cannot longer be justified in view of cellular heterogeneity - but inter-cellular molecular networks. The latter are operated by the signaling between the cytokines/chemokines and their receptors, which are readily accessible for pharmacological interventions.

## RESULTS

### Ligand-receptor genes differentially expressed between normal and cancer tissues in TCGA transcriptomes

To test the hypothesis that cancer cells are embedded in a network of “cross-talking” activity among themselves as well as with cells from the tissue microenvironment, we first curated a list of 709 ligands and 693 cognate receptors (totally 2,558 pairs of chemokine/cytokine) from various known public protein interaction databases (see details in Material and methods). We reasoned that since co-expression of a ligand and its cognate receptor is necessary (albeit not sufficient) for cell-cell communication via secreted signals, the simultaneous up- or down-regulation of expression of ligands and their cognate receptors can be the first proxy to detect gain or loss of cell-cell communication when comparing gene expression profiles from cancerous to normal tissue (Fig 1A).

**Figure 1.**
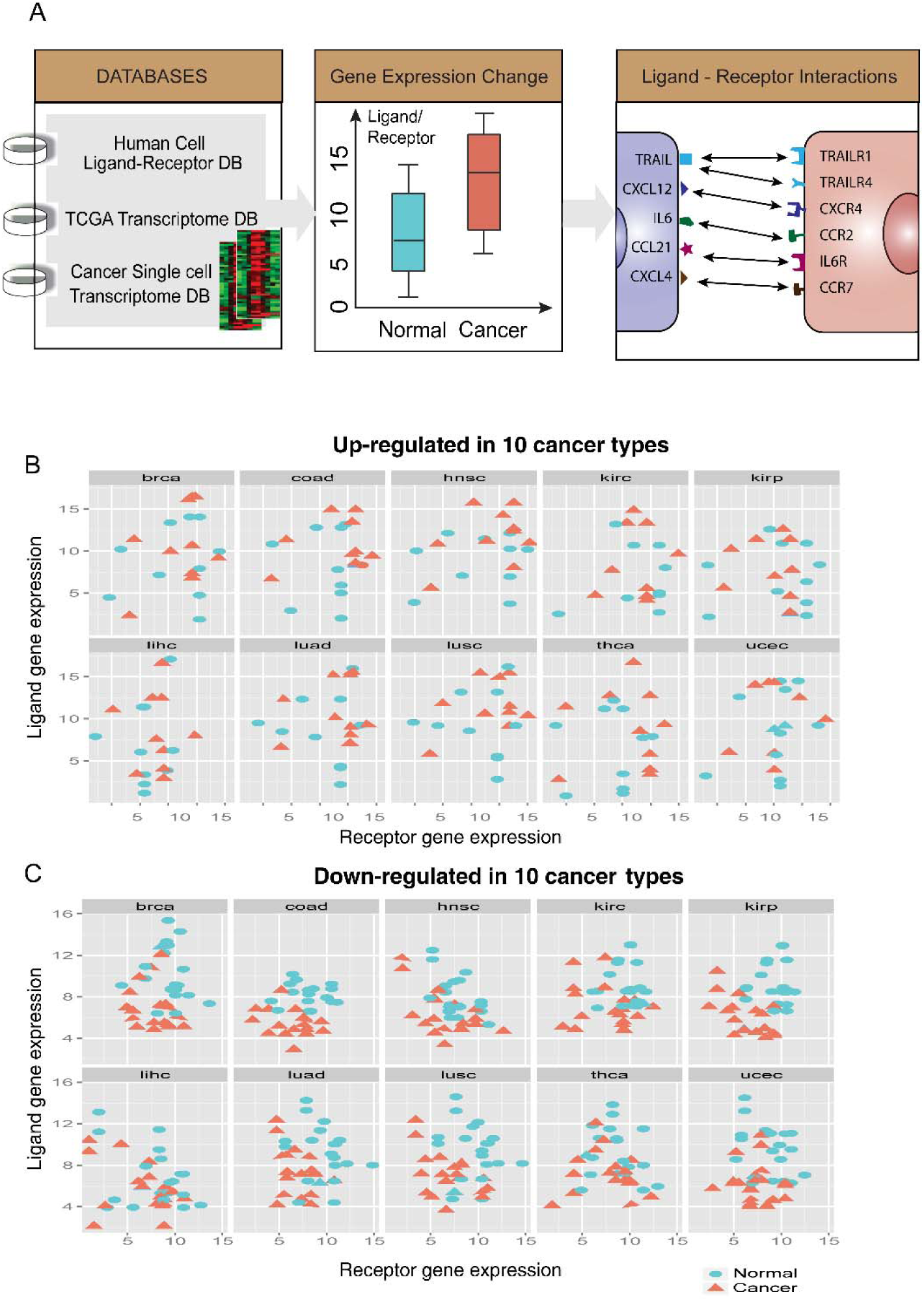
Schema & examples for the change of intercellular signaling (A) Schema to identify the changes of intercellular signaling between normal and cancer tissues from human ligand-receptor database and TCGA transcriptomes. (**B,C**) Scatter plots of top 10 ligand-receptor pairs up-regulated/down-regulated in cancer tissues compared with normal tissues. Each dot or triangle represents a population average of a ligand (y axis) and a receptor (x axis). (dot –normal tissue; triangle – cancer tissue)

We first searched for ligand-receptor pairs that were differentially expressed between the normal tissue and the cancer tissues using the TCGA database^4^, which contains a large number of patient samples. We focused on the most frequent cancer types which was necessary to achieve statistical significance for analyzing the expression changes of ligand/receptor genes in the same tumor type across the patient population. From the 30 cancer types available, we selected 10 cancers for which TCGA had at least ≥ 30 normal samples and ≥ 160 samples from distinct cancer patients. We analyzed the ligand or receptor genes that were significantly differentially expressed between the normal tissues and the cancer samples using the linear regression model (up-/down 2 folds with P-value = 0.05). The top 10 up-regulated ligand-receptor pairs across 10 cancer types are shown in Fig. 1B while the top 10 down-regulated ligand-receptor pairs are shown in Fig. 1C. To demonstrate the distribution of these most changed genes in different cancer types, boxplot of most down-regulated genes (either ligands or receptors) are shown in Fig. 2A while the most up-regulated ones are shown in Fig. 2B. For a more detailed analysis of these genes, we listed these up-/down-regulated ligand-receptor pairs which were most commonly shared across 10 cancer types for the four possible scenarios:

**Figure 2.**
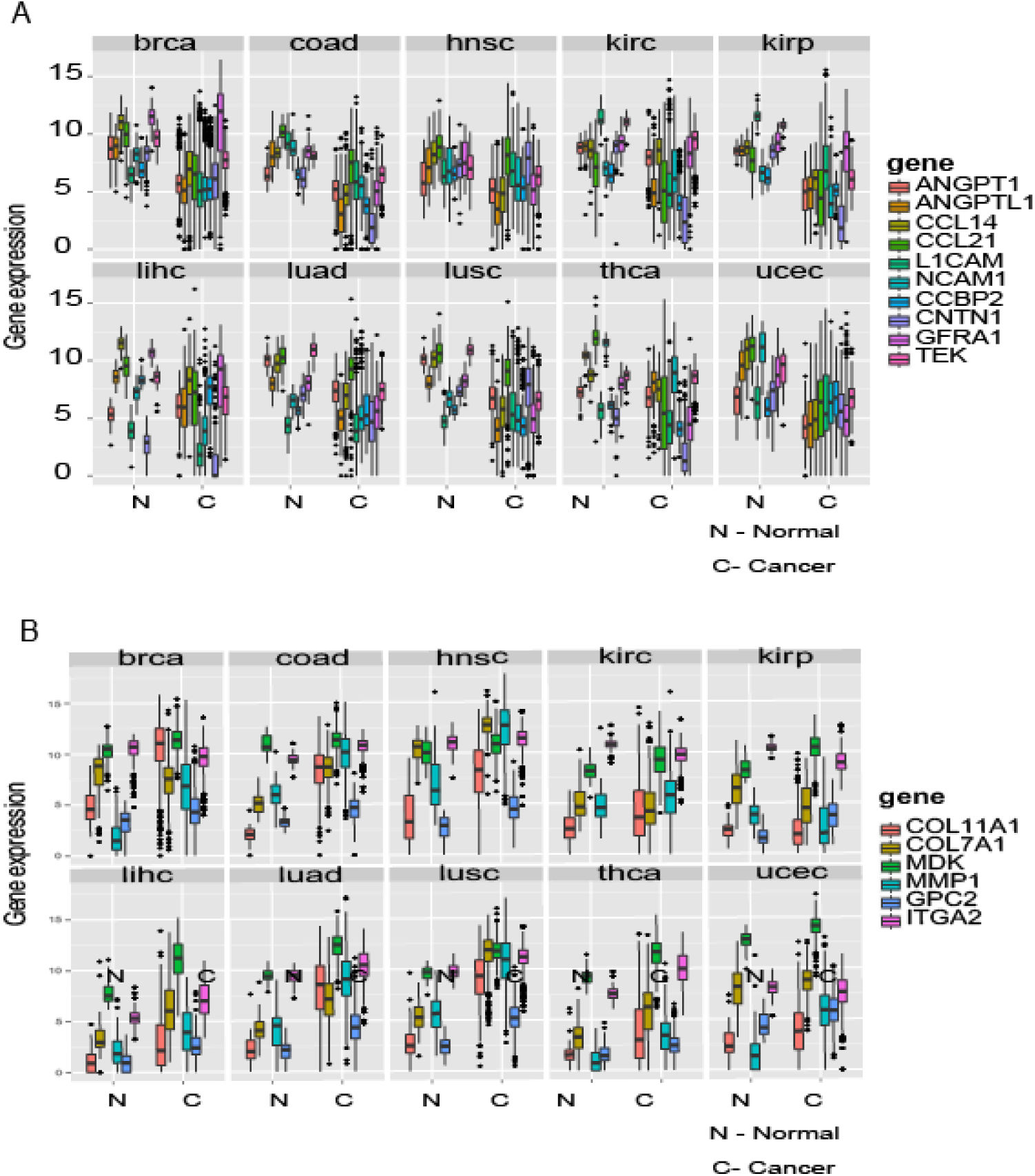
Boxplots of the most differentially expressed ligand or receptor genes across 10 cancer-normal tissue comparisons. **(A)** The boxplots of top 10 down-regulated ligand/receptor genes in cancer tissues compared with normal tissues. Each small panel is the gene expression boxplot comparison between normal (left) and cancer (right) tissues in one cancer type (brca – breast cancer etc.). Different color represents different genes. **(B)** The boxplots of top 6 up-regulated ligand/receptor genes in cancer tissues compared with normal tissues.

*(i)* Expression of both receptor and ligand increased in cancer (table 1), such as *COL11A1-ITGA2, COL7A1-ITGA2, MDK-GPC2* and *MMP1-ITGA2*. Excess expression of both Collagen *(COL11A1, COL7A1)* and Integrin *(ITGA2)* is known to be associated with cell invasiveness and tumor formation^34^. *MMP1* is an enzyme which degrades the extracellular matrix (ECM) – hence promoting cell invasiveness^35^. *MDK* is a growth factor and *GPC2* involves with *Wnt* Signaling pathway – both promote cell growth, migration and angiogenesis during tumorigenesis^36^.
*(ii)* Expression of either receptor or ligand is increased in cancer but not both (table 2), such as *CALM1-HMMR, SPP1-ITGB1, TNFSF18-TFNRSF18, COL11A1-ITGB1* etc. Not surprisingly, collagen and integrin were found also here. *CALM1-HMMR* is involved in Ca2+ controlled cell motility and invasiveness. *SPP1-ITGB1* is related to cell-matrix interactions and up-regulate stimulates the immune response. *TNFSF18-TFNRSF18* is involved with T-cell activation and lymphocytes and endothelial cell interactions.
*(iii)* Expression of both receptor and ligand genes is decreased in cancer (table 3), such as *ANGPT1-TEK, CCL21-CCBP2, CCL14-CCBP2, L1CAM-CNTN1* and *NCAM1-GFRA1*. In quiescent tissues, *ANGPT1* form complexes with *TEK* molecules to maintain cell-cell contacts, inhibiting angiogenesis and promotes the vascular quiescence. Chemokine (C-C motif) ligands *(CCL21, CCL14)* and Chemokine-binding protein *(CCBP2)* control chemokine levels and localization. *L1CAM-CNTN1* and *NCAM1-GFRA1* are both involved in cell adhesion and *NOTCH* signaling activation.
*(iv)* Expression of either ligand or receptor is decreased in cancer but not both (table 4), such as *CNTFR, LIFR, SSTR1, CXCL12, NPY1R, RELN, RSPO3, TGFBR3* etc. *CNTFR* involves with neuronal cell survival and differentiation; *LIFR* involves cellular differentiation and proliferation. *SSTR1* regulates various cellular functions such as cell proliferation and endocrine signaling – inhibiting the release of many hormones. *CXCL12* together with *CXCR4* regulates intracellular calcium ions and chemotaxis. *NPY1R* mediates diverse of biological functions, such as food intake and circadian rhythm. *RELN* is involved with extracellular matrix and regulates microtubule functions. *RSPO3* has been implicated in angiogenesis, *Wnt*/beta-catenin pathways and *TGF-beta* pathways. Decreasing expression of *TGFBR3* receptors is commonly observed in cancers.

### Altered correlation of ligand-receptor gene expression in cancer tissues in the TCGA transcriptomes

Besides considering the up-/down-regulation of ligands and their cognate receptors as proxy for cell-cell communication, correlation in the abundance of transcripts of these communication molecules across tumors would also be an indicator of cell-cell communication – irrespective of whether of the autocrine or the paracrine type. In other words, if co-expression has a biological role, one can postulate that, as a more stringent criterion for functionality, the expression change of ligands and receptors across tumors of multiple patients would be commensurate: Tumors with higher numbers of “sender” cells may also contain a larger number of “receiver” cells, which would be manifest in an increased correlation of the abundance of transcripts of ligands and receptors. Moreover, if individual cells are both sender and receiver (autocrine scenario), the varying content of such autocrine-capable cells in tumors, postulated to be a hallmark of malignancy^17^, would further increase the correlation between ligand and receptor expression across tumors.

We systematically calculated the *Spearman* correlation coefficients for the transcript abundances of the curated ligand-receptor pairs (i.e., the correlation between the ligand expression vector and the receptor expression vector across the samples for each pair) for the normal and the corresponding primary tumor tissues of all 10 cancer types. In cases where the ligands or receptors have non-bijective relationships, e.g. one ligand binds to multiple receptors, the correlation is calculated between the ligand and the sum of all receptors. The overall patterns of correlation of many ligand-receptor pairs changed significantly (Kolmogorov test for change of distributions of all the values ligand-receptor correlation coefficients): examples of ligand-receptor pairs are shown in Fig. 3. For some ligand-receptor pairs the correlation between ligands and receptors in the cancer tissue was higher, as was the case for *CCL2-CCR5, CCL3-CCR5* and *PLAU-ITGA5*. By contrast, some cancers lost the positive correlation seen in the healthy tissue, such as *LIPH-LPAR2* and *SEMA4G-PLXNB2*, and others lost the negative correlations, such as *CGN-TGFBR2* and *SEMABD-TYROBP* etc.

**Figure 3.**
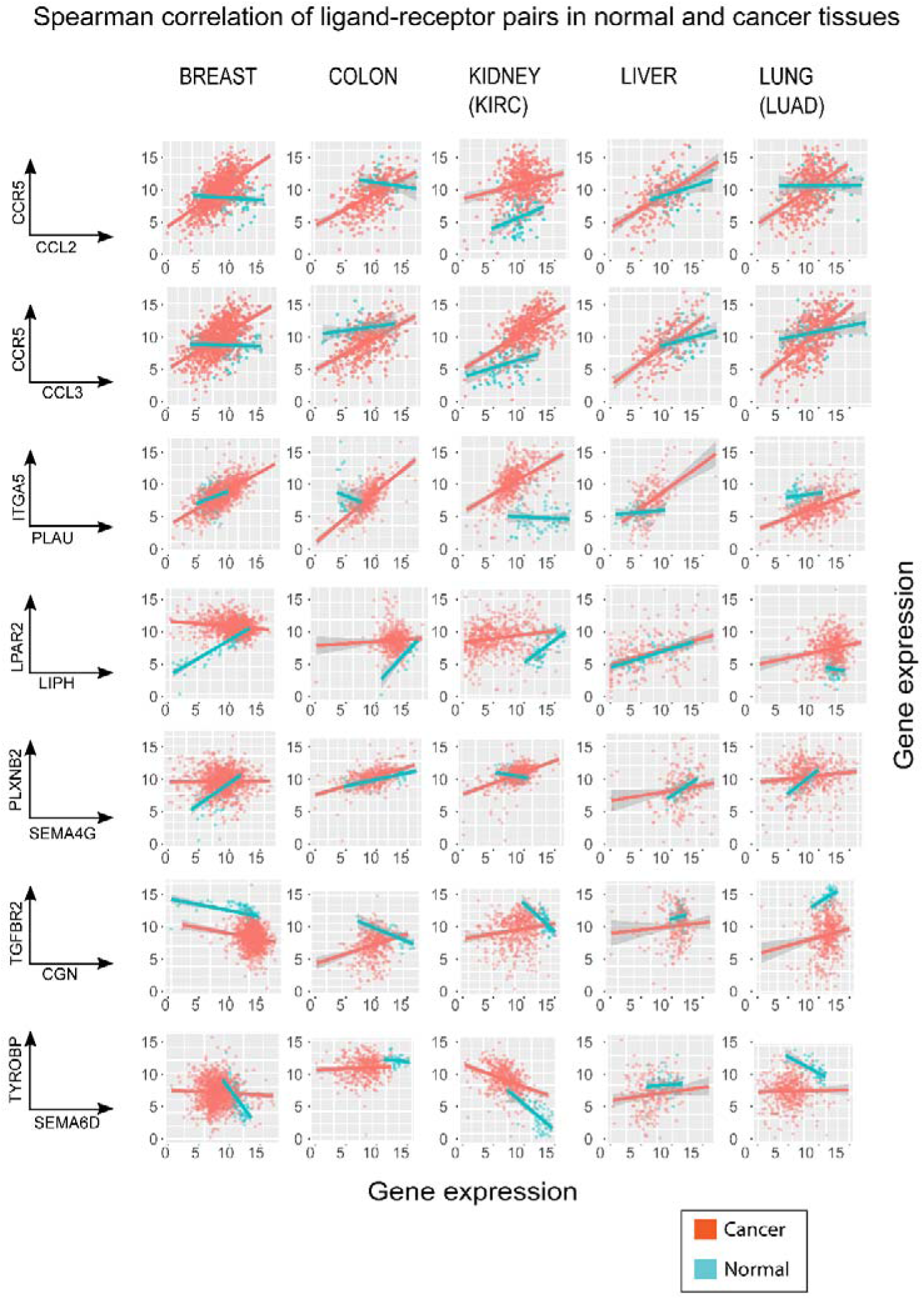
The scatter plots of seven typical ligand-receptor pairs demonstrated the altered correlations between five cancer (orange dots) and their corresponding normal tissues (blue dots). We showed the Spearman correlations of the gene expressions of ligand-receptor pairs in both normal and cancer sample for five cancer types. The ligand-receptor pairs were grouped into 3 categories: 1) gaining the correlation in cancer tissues (first 3 rows); 2) losing the positive correlation in cancer tissues (4^th^ ad 5^th^ row); 3) lose the negative correlations in cancer tissues (6^th^ and 7^th^ row). Each dot represent one ligand-receptor gene expression in one sample. The line is a regression line to visualize the trend of the data.

To obtain a comprehensive view of ligand-receptor correlation of all curated pairs, we plotted the histograms of the values of the ligand-receptor *Spearman correlation coefficients* for all pairs in normal and cancers tissues for the 10 cancer types (as shown in Fig. 4A). This aggregate analysis revealed that the spread of the distribution of ligand-receptor Spearman correlation coefficients consistently shrank: tumors of most cancer types lost both the high values of positive and negative correlation. This change of the distribution of correlation coefficients between cancer and normal tissues was significant (Kolmogorov-Smirnov test, P-value < 0.01, see Table 5 and details in Materials and methods). Fig 4B shows the ligand/ receptor pairs with the most altered correlation detected by Spearman correlation.

**Figure 4.**
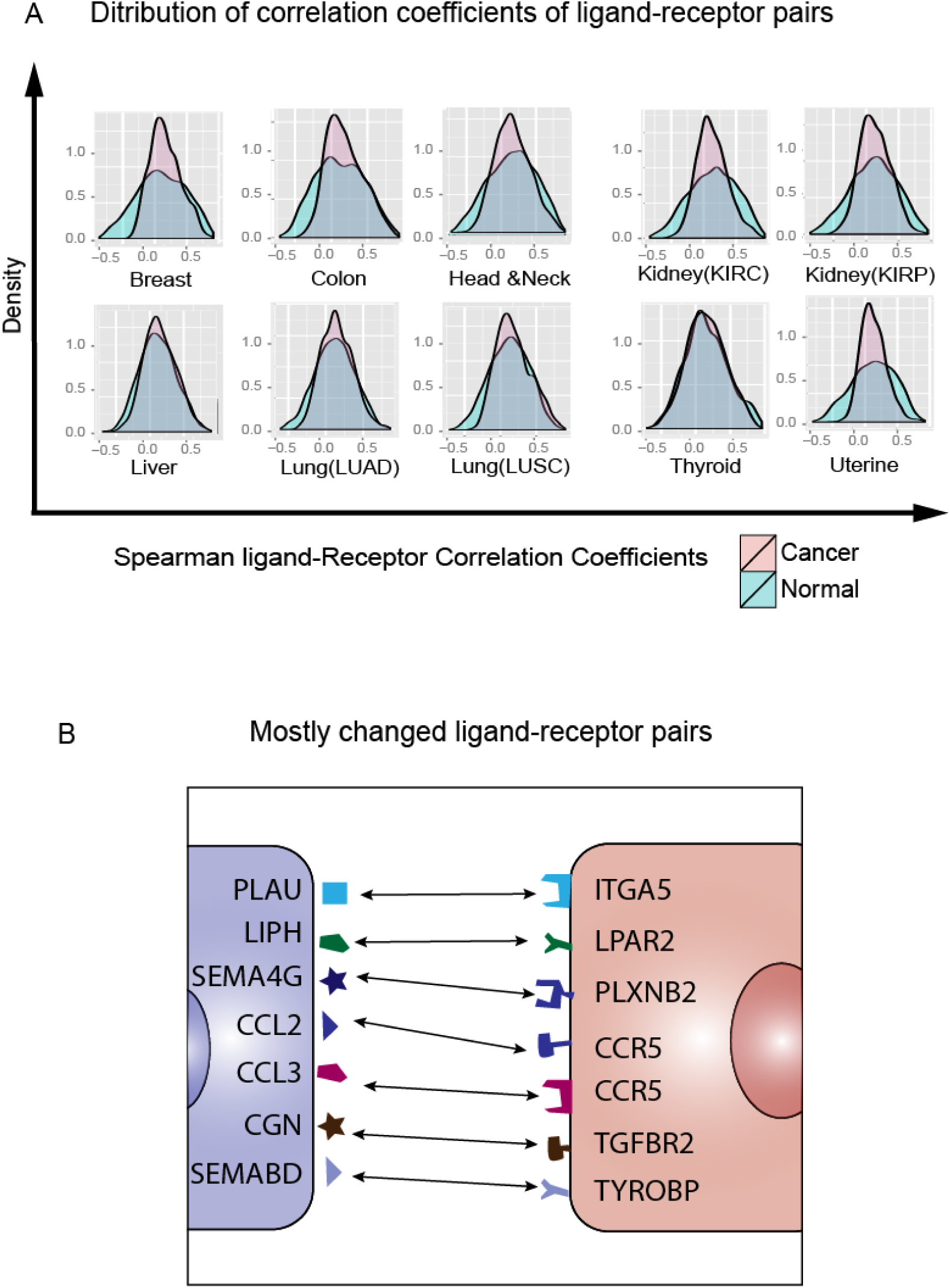
The overall Spearman correlation distribution between ten normal and cancer tissues. (**A**) The histograms of Spearman correlation coefficients of 2,558 ligand-receptor pairs between 10 cancer samples (in *red*) and their corresponding normal tissue samples (in *blue*). The ligand-receptor pairs lose the correlation in most cancer tissues. (**B**) The ligand-receptor pairs shared in most cancers with altered correlations between 10 cancer and normal tissues.

As negative control for the significance of the correlation in the expression of ligands and their cognate receptors, we computed the correlation of random pairs of receptors and ligands (which do not interact specifically). The ligand-receptor correlation for all possible random pairs (2,558 × (2,558-1) / 2 = 3,270,403 pairs) of ligands and receptors, irrespective of their binding specificity, was computed to serve as background signal. As shown in Fig. 5A, the distribution of correlation coefficients of ligand-receptor pairs of cancer tissues extracted from the TCGA gene expression data have a much higher peak around zero than the one derived from the random pairs, indicating that overall, specific ligand-receptor pairs exhibit lower variability of correlation than one would expect from random pairs, warranting the use of ligand-receptor correlation as an indicator of a biological change in the tumor tissue. For most cancer types (breast, head and neck, liver, lung (LUSC) and thyroid), the distribution had a fat long tail of higher correlation values for specific ligand/receptor pairs compared to random pairs – in line with the small subset of ligand-receptor pairs that displayed a significantly higher correlation in cancer tissues. Note that the differences of the fat tails do not exist in the comparison between observed ligand-receptor pairs and random pairs of the normal tissues (Fig. 5B).

**Figure 5.**
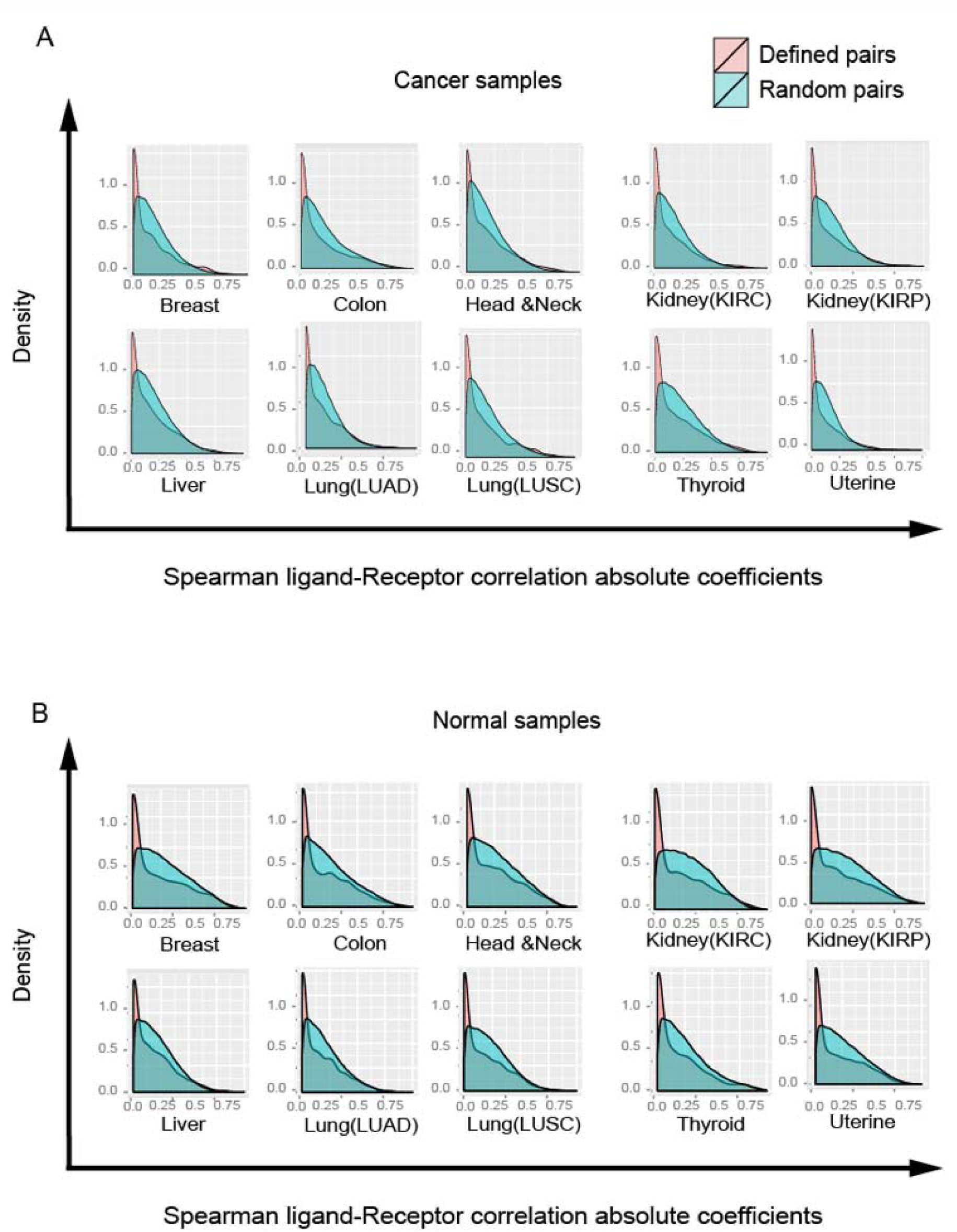
Spearman correlations in cancer and normal tissues are significantly different from randomly chosen pairs. **(A)** The Spearman correlation coefficients distribution (random (in red color) vs. defined (in blue) ligand-receptor gene pair) in cancer tissues. **(B)** The Spearman correlation coefficients distribution (random vs. defined ligand-receptor gene pair) in normal tissues.

Next, to display the nature of the changes, we plotted the values of *Spearman* correlation coefficients of the 2,558 ligand/receptor pairs as scatter plots for a cancer (Y-axis) against the corresponding normal tissue (X axis) (Fig. 6). The ligand-receptor pairs whose relationship of expression in cancer differed most from that in normal tissue could be divided in three categories, indicated by the three rectangle boxes in the scatter plots away from the diagonal: *(i)* uncorrelated in normal (*Spearman* correlation coefficients between -0.25 and 0.25) but correlated in cancer (higher than 0.5), located in rectangle area *I* in the scatter plot of Fig. 6C; (*ii*) negatively correlated in normal (lower than -0.5) but uncorrelated (between -0.25 and 0.25), in rectangle area *II* of Fig. 6C and *(iii)* positively correlated in normal (higher than 0.5) but uncorrelated (between −0.25 and 0.25) in cancer, located in rectangle area *III* (Fig. 6C). In order to estimate the background noise, we had randomized two groups of 65 samples from the 112 samples of normal breast tissue and compared their fluctuations in the scatter plots (Fig. 6A and 6B). This indicates that variations of ±0.25 from the value of correlation coefficients were within the range of measurement noise; hence we defined as ‘uncorrelated’ *Spearman* correlation coefficients between -0.25 and 0.25, and as ‘positively’ or ‘negatively correlated’ values higher than 0.5 or lower than -0.5, respectively. With these criteria we systematically compared ligand-receptor correlation differences through all scatter plots for all ten cancer tissues (Fig. 6C – 6O).

**Figure 6.**
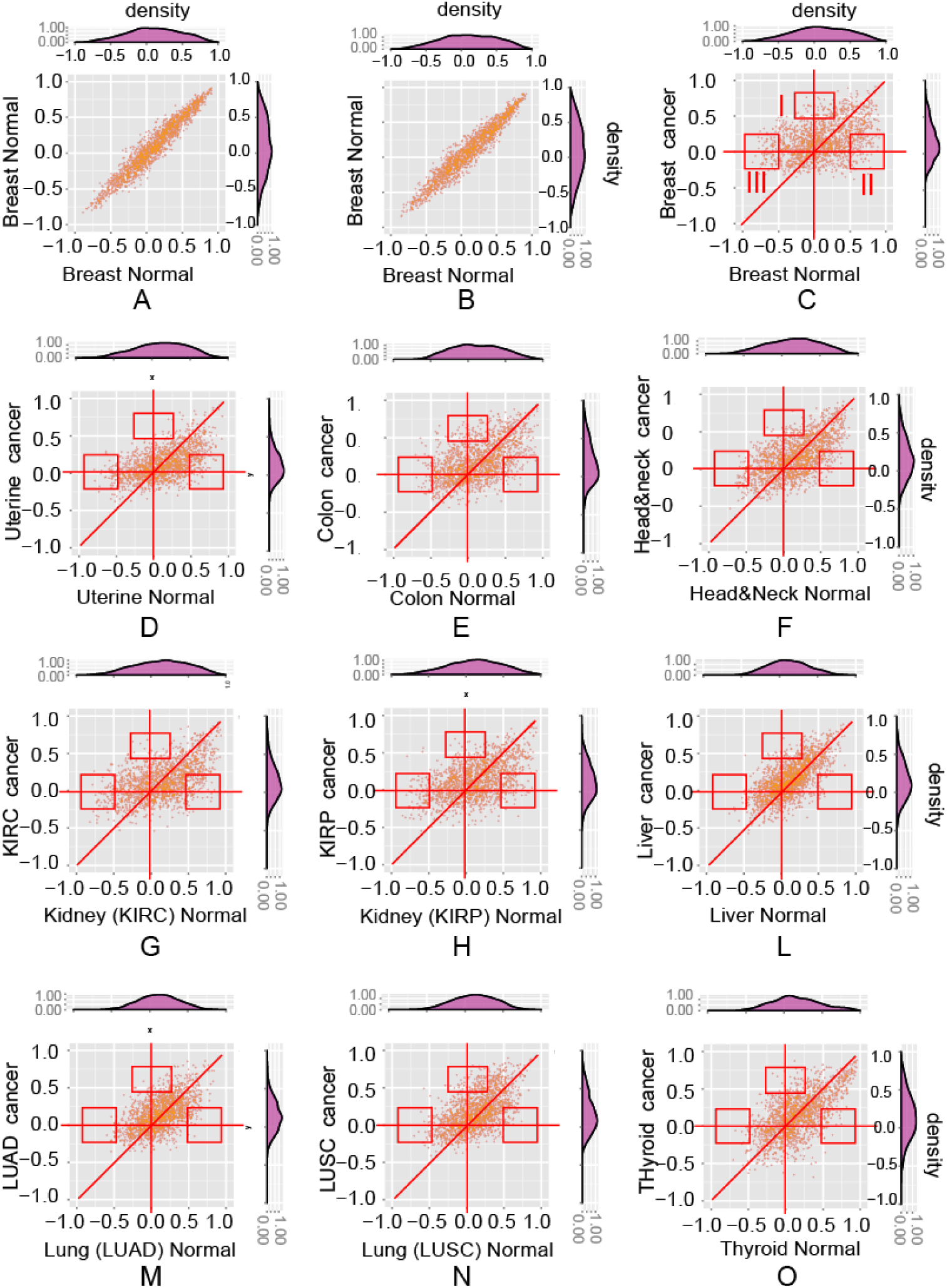
The scatter plots show the altered ligand-receptor correlations between normal and cancer tissues of 10 cancer types. **(A)(B)**. the background noise of correlations in the normal breast tissue. The scatter plots of 2,558 ligand-receptor pairs from 65 samples which are randomly chosen 1,000 times from breast tissue (performed twice); **(C)-(O)**. The scatter plots of ligand-receptor correlations (normal vs. cancer) of ten cancer types. Three types of altered correlations: i) area *I*: changing from uncorrelated (Spearman correlation coefficients between - 0.25 and 0.25) to correlated (higher than 0.5); ii) area *II*: changing from negatively correlated (lower than -0.5) to uncorrelated (between −0.25 and 0.25); iii) area *III*: changing from positively correlated (higher than 0.5) to uncorrelated (between -0.25 and 0.25).

Among the several pairs identified in this survey, some have been implicated in cancer initiation and progression, often impacting angiogenetic processes, immune responses facilitating interactions between cancer cells and carcinoma associated fibroblasts. This was, for instance, the case with *CCR5* which, with the two ligands i.e. *CCL2* and *CCL3*, formed the two ligand-receptors pairs that exhibited the largest increase in correlation when comparing tumor to normal tissues (Table 2). *CCR5* is expressed on several tumor-associated immune cells, including T cells, dendritic cells, and NK cells as well as endothelial and cancer cells. *CCR5* and its ligands *CCL2, CCL3, CCL4* and *CCL8* play crucial role in migration, activation and proliferation of T-cells and polarization of macrophages. It is also involved in metastasis and mediates the recruitment of endothelial cells, thus contributing to angiogenesis^37,38^. Furthermore, the *CCR5-CCL4* axis can contribute to breast cancer metastasis to bone by mediating the interaction between cancer cells and fibroblasts in bone cavity^39^. Another well-known pair is the *CCL2-CCR2* axis which mediates macrophage recruitment, promotes tumor growth, progression and metastases in breast and prostate cancers^40,41^. Furthermore, evidence suggests that carcinoma associated fibroblasts (CAFs) recruit monocytes mostly through the *CCL2-CCR2* axis in breast and melanoma cancers^42,43^. In a mouse model of pancreatic cancer, *CCR2* inhibitors depleted inflammatory monocytes and macrophages, which resulted in decreased tumor growth and reduced metastasis^44^. It is well known that *CCL2* promotes lymphocyte, keratinocyte, and endothelial cell activation; furthermore, *CCL2* can indirectly affect angiogenesis. The dysregulation of chemokines that activate neutrophils or monocytes / macrophages can negatively impact angiogenesis by limiting the secretion of proangiogenic factors^45^. The top 15 ligand-receptor pairs for which the ten cancer types showed most significant alteration in ligand-receptor correlation in three categories are listed in Tables 6 to Table 8 separately.

### Role of immune cell infiltration in modulating the ligand-receptor correlations

The altered expression and correlation of ligand-receptor pairs in tumors most likely originates from the altered interaction patterns of tumor cells with the infiltrating immune cells. Inflammatory cells in the tumor stroma interact with tumor cells in a bi-directional way via cytokines, such as *IL6, IL8* and many others, which can promote tumor progression or mediate an immune response to the neoplastic cells^21^. Analysis of gene expression profiles in cancer cell lines^46,47^ as well as measurements in cancer cell culture conditioned media^48,49^ suggest that these cytokines, originally associated with immune cells, can also be produced by carcinoma cells.

Thus, we next sought to assess the contribution of immune cell infiltrates to the altered correlation of ligand-receptors expression in tumors. We took advantage of the information in TCGA that offers an estimate of the immune and other stromal cell contents in each tumor sample. This had previously been calculated by Carter et.al. from somatic DNA alternations in exome sequencing data^50^. We divided the primary tumor samples into two groups: tumors in which the proportion of infiltrating non-neoplastic cells was *(i)* high 25% quantile or *(ii)* low 25% quantile. This distinction was done with respect to four types of non-neoplastic admixture cells: lymphocyte, monocytes, neutrophil and stromal cells. If these micro-environmental cells contribute substantially to cell-cell communication, that is, to the increase in tumors of the correlation of expression between ligand and receptor, the correlation coefficient should be significantly different in between tumors of these two groups that have very different nonneoplastic cell content. Fig. 7 shows that the ligand-receptor correlation coefficients was not different between tumor samples with high or low content of stromal infiltrates, because PCA analysis cannot separate these two groups as two clusters in the first two principal components. This result indirectly suggests that scaling of transcript expression of ligands and their cognate receptors in tumors, which indicate cell-cell communication, is not simply a manifestation of increased interaction between cells of the immune infiltrate and the tumor parenchyma.

**Figure 7.**
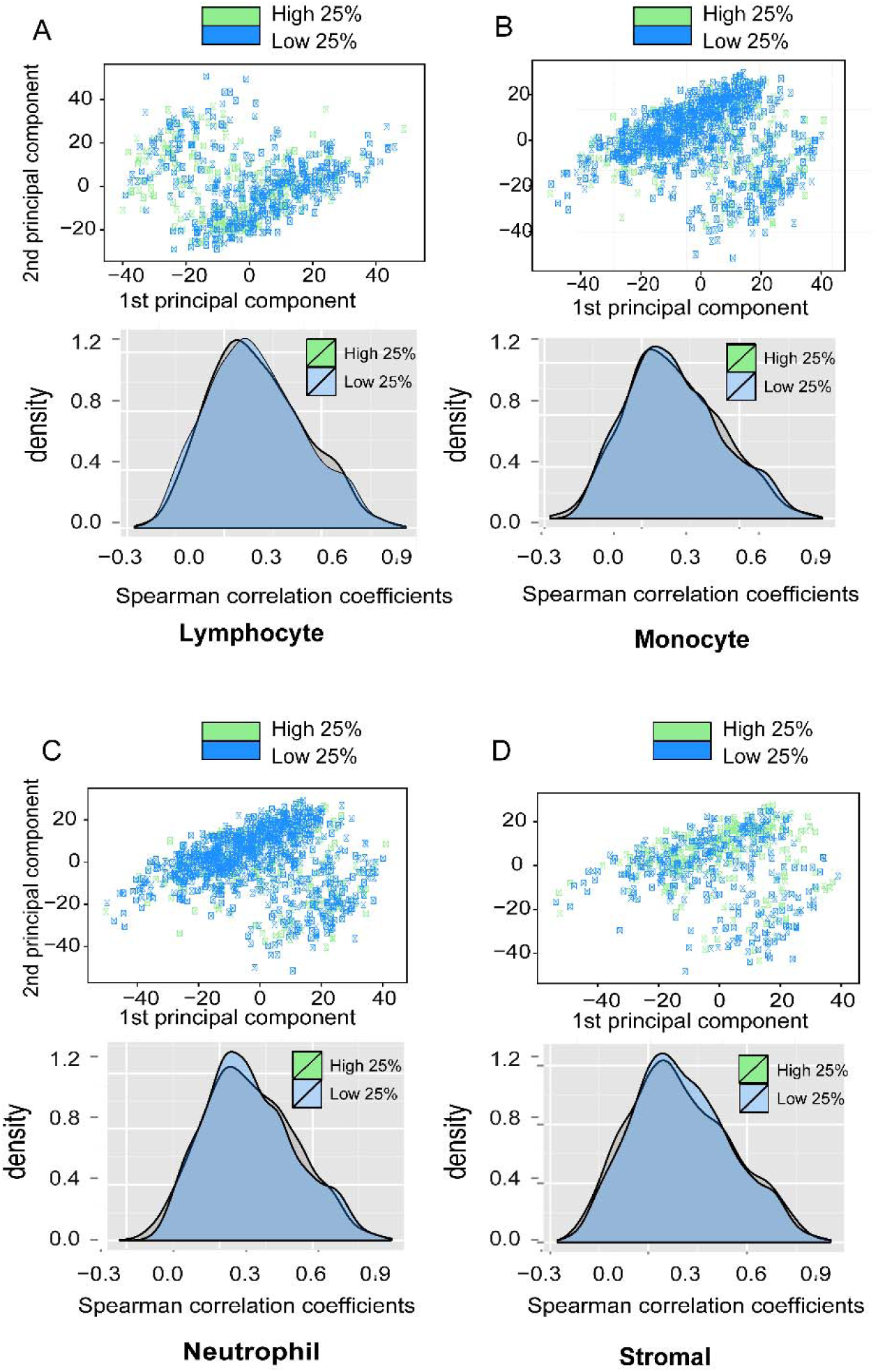
Spearman correlation coefficient distributions of ligand-receptor pairs of breast cancer tissues with different percentage of immune cells. The scatter plots in first 2 PCA components and histograms of ligand-receptor correlation distributions of two groups of tumor samples: top 25% high inflammation cell infiltration group (in green) vs. bottom 25% low inflammation cell infiltration group (in blue). **(A)** Lymphocyte **(B)** monocyte **(C)** Neutrophil **(D)** Stromal cells.

### Ligand-receptor co-expression analysis in single-cell gene expression data

The arrival of single-cell transcriptome data now offers the opportunity for mapping a finegrained cell-cell communication scheme for how the receiving (receptor-expressing) and signaling (ligand expressing) cells are distributed among the various cell subpopulations of cells in the tumor. Single cell-transcriptomics of the mixed population of dissociated cells from a tumor readily identifies the cell types in addition to providing information on expression of ligands and receptors by individual cells, and thus allows us to determine “who talks to whom” in the cell-cell communication network. We demonstrate this using one of the first data sets available as an example for future analyses as such single-cell resolution data becomes available for many tumor types. Garraway et. al. published a set of single cell-RNASeq data that revealed intratumoral heterogeneity of gene expression in primary and metastatic melanoma^51^. We analyzed the same 2,558 ligand-receptor pairs in the transcriptomes of ~4,000 individual cells isolated from melanoma which include both malignant tumor cells and non-tumor stroma and immune cells. We systematically compared the ligand-receptor expression patterns among seven cell types present in the melanoma-derived single-cell suspension (Melanoma cells, T cells, B cells, macrophage, NK cells, CAF and endothelial cells)^51^. We then built the interaction network of indicating which cell types “talks” to which other cell type, based on the specific ligand-receptor link. The gene expression distribution in single-cell transcriptomes obtained from RNASeq is usually modeled as a negative binomial distribution^52^ If the gene expression values *x_i_* of both the ligand and the receptor were above a certain threshold for two individual cells, this would indicate a possible intercellular signaling between these two cells. We used the following criteria to call a cell-cell dialogue between two different cells as existent:

We calculated the mean 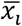 and standard deviation *σ_i_* of the expression level of a gene *i* (either ligand or receptor) across all seven cell subpopulations representing the seven cell types. If a subpopulation’s average gene expression level was larger than 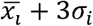, it was considered to be highly expressed in that subpopulation. Where both members of a ligand-receptor pair were highly expressed in two cell types out of the seven cell subpopulations, we interpreted this as a potential cell-cell communication line between these two cell types. The above threshold is tunable and we used here the above value for demonstration purposes. A hence identified ligand-receptor pair also indicates the direction of a communication line (sender→receiver), allowing us to draw a communication network as a directed graph, as show in Fig 8A.

**Figure 8.**
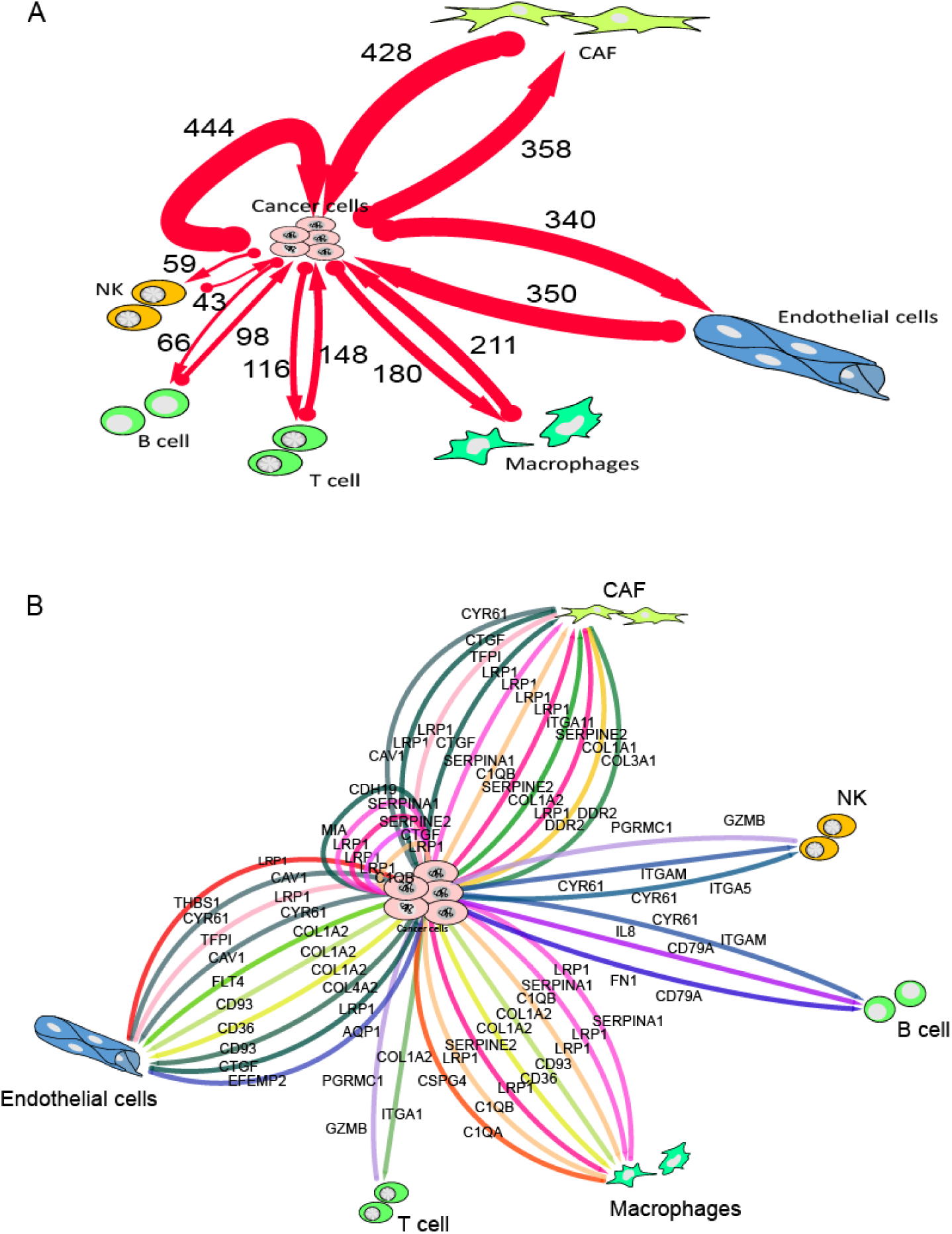
The proposed cell-cell interaction network between cancer and the normal cells in the microenvironment derived from melanoma single-cell data. To highlight the crosstalk between cancer cells and the microenvironment, we filtered out all the interactions not involved in cancer cells. (**A**) The network shows the number of all the proposed ligand-receptor interactions involving melanoma cancer cells and the normal cells in the micro-environment. The majority of the interactions seem to involve the endothelial cells, CAFs and cancer cells themselves. (**B**) The top 2% of the ligand-receptor pairs cross-talking between cancer cells and normal cells. The ranking is based on the exclusiveness score of the interaction (exclusiveness score: gene expression level of one cell type divided by the mean across all cell types).

Using this analysis framework we found that by far most communication lines occurred between cancer and endothelial cells, and between cancer cells and cancer associated fibroblasts (CAF), as well as among the cancer cells themselves. Using this analysis, we could also label the links in the cell-cell interaction network map in the melanoma tissues with the specific by ligand-receptor pairs (Fig. 8B): melanoma tumor cells talked to CAF cells mainly through the three ligand-receptor pairs *CTGF-LRP1, C1QB-LRP1* and *COL1A2-ITGA11;* conversely, CAF cells “talked back” through *CYR61-CAV1, CTGF-LRP1* and *COL3A1-DDR1*. Tumor cells talked to macrophages through the ligand receptor pairs *C1QB-LRP1, COL1A2-LRP1* and *SERPINE2-CD36*, and macrophages talked back through *SERPINA-LRP1, C1QB-LRP1* and *C1QA-CSPG4;* tumor cells talked to T-cells through *COL1A2-ITGA1* and T cells talked back through *GZMB-PGRMC1*. Tumor cells talked to endothelial through the pairs *COL1A2-FLT4, COL1A2-CD93* and *COL4A2-CD93;* and endothelial cells talked back through *THBS1-LRP1, CYR61-CAV1* and *TGPI-LRP1*. Finally, tumor cells talked to NK cells through *CYR61-ITGAM* and *CYR61-ITGA5* and NK cells talked back through *GZMB-PGRMC1*. As can be seen from this list,many interactions where mediated by collagen deposition, acting as a ligand to a variety of receptors.

## DISCUSSION

In recent years many reports, such as the discovery of cancer without driver mutations^53^ and collective metastasis^54^ etc., have challenged the traditional paradigm of tumorigenesis in which cancer cells accumulate mutations in genes that control growth, which lead to clonal expansions and metastasis at late stage^55^. Among the recent explanations proposed to address these inconsistencies with the oncogene paradigm, a more comprehensive and dynamic view describes tumors as complex tissues consisting of multiple different cell types that communicate with each other^9,11^. Thus, cancer is an emergent system: tumor growth and progression are promoted by the intricate network of cooperation among cancer cells themselves and between cancer cells and (non-neoplastic) cells and matrix of the tumor micro-environmt.

The emphasis on cellular interactions among cancer and normal cell populations is based on several findings and one of the most convincing is the presence of a large array of growth factors, chemokine/cytokines and matrix proteins detected in tumor tissues. Furthermore, many of these secreted molecules are produced not only by the non-neoplastic stromal tissues but also by the cancer cells themselves and directed at other cancer cells. Since it is expensive and time-consuming to directly measure the ligand-receptor interactions in hundreds of tumor samples, the up/down-regulation and the *Spearman correlation* of ligand-receptor pair was used as the surrogate of the existence of the ligand receptor interaction as well. If a ligand interacts with a receptor, depending on their molecular structure, one receptor could bind to one, two or even more ligands – certain quantitative relationship can exist between the ligand and the receptor. Our analysis revealed the pronounced changes in gene expression or correlation of several cyto-chemokines/receptors pairs in most cancer tissues. Among the most frequent pairs of chemokines/receptors that whose transcript expression was upregulated or the correlations increased in tumors were: *COL11A1-ITGA2, COL7A1-ITGA2* and *MMP1-ITGA2*, likewise *CCR5* and the ligands *CCL2, CCL3* etc. All these receptors/ligands molecules are endowed with several tumorigenic properties, such as collagen-integrin interaction, cell-ECM(extracellular matrix) interactions and Ca2+ controlled cell-ECM interactions. The most frequently down-regulated pair of chemokines/receptors were *ANGPT1-TEK, CCL21-CCBP2, CCL14-CCBP2, L1CAM-CNTN1* and *NCAM1-GFRA1 etc*. Their biological functions are to inhibit angiogenesis, maintain vascular quiescence and cell adhesion, modulate immune-regulatory and inflammatory processes etc.

We calculate the Spearman correlation to detect such quantitative relationship – the higher the correlation, the more possibly quantitative relationship exist between the ligand-receptor pair. However, two issues may complicate the analysis. 1) most ligand receptor gene expression data are quite noisy (Spearman correlation coefficients mostly vary from -0.3 to 1 in cancer tissues), thus the correlative measurements presented here represents only a first step, broadly applied step towards identifying receptor ligand airs involved in tumorigenesis. We need a better measure to represent the ligand receptor interaction; 2) even we analyzed the influence of immune cells and found no obvious influence, it needs a well-thought experiment to distinguish the changes in ligand-receptor quantitative relationship which are caused by cancer cell interactions or the infiltration by immune cells.

The cell-cell communication network we calculated from single-cell melanoma relies on the specificity of the interaction between a given ligand produced by a given cell and the its cognate receptor expressed by a different cell, on the large variety of cancer and stromal cells and on the expression profiles which define which ligand and which receptor they express. Future availability of single-cell transcriptomes will expand such analysis to other tumor cell types, affording a new observable in the characterization of tumor phenotype: the structure of the cell-cell communication network – which may be used for comparison, classification and even as prognostic and predictive marker of cancer.

## METHODS

### Curated ligand-receptor Database

We used the database provided by Ramilowski et.al.^56^ which includes 2,558 unique human ligand-receptor interactions. They firstly merged the lists of ligand-receptor pairs from Database of Ligand-Receptor Partners (DLRP)^17^, IUPHAR^57^ and Human Plasma Membrane Receptome (HPMR)^58^, then further extended by inferring experimentally supported interactions in the HPRD and STRING database.

### Ten solid tumor transcriptomes from TCGA

Gene expression data were retrieved from The Caner Genome Atlas (TCGA) project data portal (https://tcga-data.nci.nih.gov/). We focused our analysis on ten tissues: breast invasive carcinoma (BRCA), Uterine Corpus Endometrial Carcinoma (UCEC), Colon adenocarcinoma (COAD), Head and Neck squamous cell carcinoma (HNSC), Kidney renal clear cell carcinoma (KIRC), Kidney renal papillary cell carcinoma (KIRP), Liver hepatocellular carcinoma (LIHC), Lung adenocarcinoma (LUAD), Lung squamous cell carcinoma (LUSC), Thyroid carcinoma (THCA). We considered the samples annotated as 01 to be “cancer” (primary solid tumor) and the ones annotated as 11 to be “normal” (solid tissue normal).

### Calculate correlation of ligand-Receptor pairs

For each pair of ligand and receptor we calculated the *Spearman* correlation coefficient. We used the *Spearman* correlation function from the base package of R version 3.3.1. Three different groups of interactions were defined based on the value of their correlation. The group that gain positive correlation in cancer: we focused on the pairs that have a coefficient > 0.5 in cancer and a coefficient > -0.25 & < 0.25 in healthy tissues; the group that lose positive correlation in cancer: we focused on the pairs that have a coefficient > 0.5 in the healthy tissues and a coefficient > -0.25 & < 0.25 in cancer. The group that lose negative correlation in cancer: we focused on the pairs that have a coefficient < -0.5 in healthy tissues and a coefficient > -0.25 & < 0.25 in cancer. Once the pairs in each group defined, we focused our attention on the most recurrent ligand-receptor pairs across the ten cancer types analyzed.

### Kolmogorov-Smirnov test of the significance of correlation coefficients distribution differences between cancer and normal tissue

We use the Kolmogrov-Sminov test to calculate the distances and P-values of *Spearman* correlation coefficients distribution between 10 normal and cancer tissues from TCGA. The distribution is estimated from the numeric sets of correlation coefficients. The null hypothesis to calculate P-values is that we draw two datasets from the same continuous distribution.

## Acknowledgements

We thank Thea Tlsty (UCSF) for advice on the cancer-microenvironment interactions. This work was supported by National Institute of General Medical Sciences (NIGMS) Grant R01GM109964 and NIGMS National Centers for Systems Biology Grant 2P50GM076547-06A1.

## Author Contributions Statement

J.X.Z., R.T. and E.P. curated data and performed data analysis. S.H. conceived the idea and the theoretical analysis. J.X.Z, R.T. and S.H. drafted the manuscript, E.P. and T.K. edited and wrote the paper.

## Data availability statement

The transcriptome data of 10 cancer types from TCGA and single-cell RNASeq data of melanoma are publicly available.

